# Whole genome sequences of Malawi cichlids reveal multiple radiations interconnected by gene flow

**DOI:** 10.1101/143859

**Authors:** Milan Malinsky, Hannes Svardal, Alexandra M. Tyers, Eric A. Miska, Martin J. Genner, George F. Turner, Richard Durbin

**Author notes:** Correspondence to (MM), (RD). These authors contributed equally to this work (Milan Malinsky, Hannes Svardal).

## Abstract

The hundreds of cichlid fish species in Lake Malawi constitute the most extensive recent vertebrate adaptive radiation. Here we characterize its genomic diversity by sequencing 134 individuals covering 73 species across all major lineages. Average sequence divergence between species pairs is only 0.1-0.25%. These divergence values overlap diversity within species, with 82% of heterozygosity shared between species. Phylogenetic analyses suggest that diversification initially proceeded by serial branching from a generalist *Astatotilapia-like* ancestor. However, no single species tree adequately represents all species relationships, with evidence for substantial gene flow at multiple times. Common signatures of selection on visual and oxygen transport genes shared by distantly related deep water species point to both adaptive introgression and independent selection. These findings enhance our understanding of genomic processes underlying rapid species diversification, and provide a platform for future genetic analysis of the Malawi radiation.

**One Sentence Summary**: The genomes of 73 cichlid fish species from Lake Malawi uncover evolutionary processes underlying a large adaptive evolutionary radiation.

## Main Text

The formation of every lake or island represents a fresh opportunity for colonization, proliferation and diversification of living forms. In some cases, the ecological opportunities presented by underutilized habitats facilitate adaptive radiation - rapid and extensive diversification of the descendants of the colonizing lineages^1–3^. Adaptive radiations are thus exquisite examples of the power of natural selection, as seen for example in Darwin’s finches in the Galapagos^4,5^, Anolis lizards of the Caribbean^6^ and in East African cichlid fishes^7,8^.

Cichlids are one of the most species-rich and diverse families of vertebrates, and nowhere are their radiations more spectacular than in the Great Lakes of East Africa: Malawi, Tanganyika, and Victoria^2^, each of which contains several hundred endemic species, with the largest number in Lake Malawi^9^. Molecular genetic studies have made major contributions to reconstructing the evolutionary histories of these adaptive radiations, especially in terms of the relationships between the lakes^10,11^, between some major lineages in Lake Tanganyika^12^, and in describing the role of hybridization in the origins of the Lake Victoria radiation^13^. However, the task of reconstructing within-lake relationships and of identifying sister species remains challenging due both to retention of large amounts of ancestral genetic polymorphism (i.e. incomplete lineage sorting) and potential gene flow between taxa^12,14–18^.

Initial genome assemblies of cichlids from East Africa suggest that an increased rate of gene duplication together with accelerated evolution of some regulatory elements and protein coding genes may have contributed to the radiations^11^. However, understanding of the genomic mechanisms contributing to adaptive radiations is still in its infancy^3^. Here we provide an overview of and insights into the genomic signatures of the haplochromine cichlid radiation of Lake Malawi.

The species that comprise the Lake Malawi haplochromine cichlid radiation can be divided into seven groups with differing ecology and morphology (see Supplementary Note): 1) the rock-dwelling ‘mbuna’; 2) *Rhamphochromis* - typically midwater pelagic piscivores; 3) *Diplotaxodon*-typically deep-water pelagic zooplanktivores and piscivores; 4) deep-water and twilight feeding benthic species; 5) ‘utaka’ feeding on zooplankton in the water column but breeding on or near the lake bottom (here utaka corresponds to the genus *Copadichromis);* 6) a diverse group of benthic species, mainly found in shallow non-rocky habitats. In addition, *Astatotilapia calliptera* is a closely related generalist that inhabits shallow weedy margins of Lake Malawi, and other lakes and rivers in the catchment, as well as river systems to the east and south of the Lake Malawi catchment. This division into seven groups has been partially supported by previous molecular phylogenies based on mtDNA and amplified fragment length polymorphism (AFLP) data^18–20^. However, there have also been substantial differences between the published phylogenies and, in particular, the question of whether the groups are genetically separate remained unanswered.

To characterize the genetic diversity, species relationships, and signatures of selection across the whole radiation, we obtained paired-end Illumina whole-genome sequence data from 134 individuals of 73 species distributed broadly across the seven groups (Fig. 1a; see Supplementary Note). This includes 102 individuals at ~15× coverage and 32 additional individuals at ~6× (Supplementary Table 1).

### Low genetic diversity and species divergence

Sequence data were aligned to the *Metriaclima zebra* reference assembly version 1.1^11^, followed by variant calling restricted to the ‘accessible genome’ (the portion of the genome where variants can be determined confidently with short read alignment), which comprised 653Mb or 91.5% of the assembly excluding gaps (see Supplementary Methods). Average divergence from the reference per sample was 0.19% to 0.27% (Supplementary Fig. 1). Across all samples, after filtering and variant refinement, we obtained 30.6 million variants of which 27.1 million were single nucleotide polymorphisms (SNPs) and the rest were short insertions and deletions (indels). Unless otherwise indicated, the following analyses are based on biallelic SNPs.

To estimate nucleotide diversity (π) within the sampled species we measured the frequency of heterozygous sites in each individual. The estimates are distributed within a relatively narrow range between 0.7 and 1.8×10^−3^ per bp (Fig. 1b). The mean π estimate of 1.2×10^−3^ per bp is at the low end of values found in other animals^21^, and there appears to be little relationship between π and the rate of speciation: individuals in the species-rich mbuna and shallow benthic groups show levels of π comparable to the relatively species-poor utaka, *Diplotaxodon*, and *Rhamphochromis* (Supplementary Fig. 1).

**Fig. 1:**
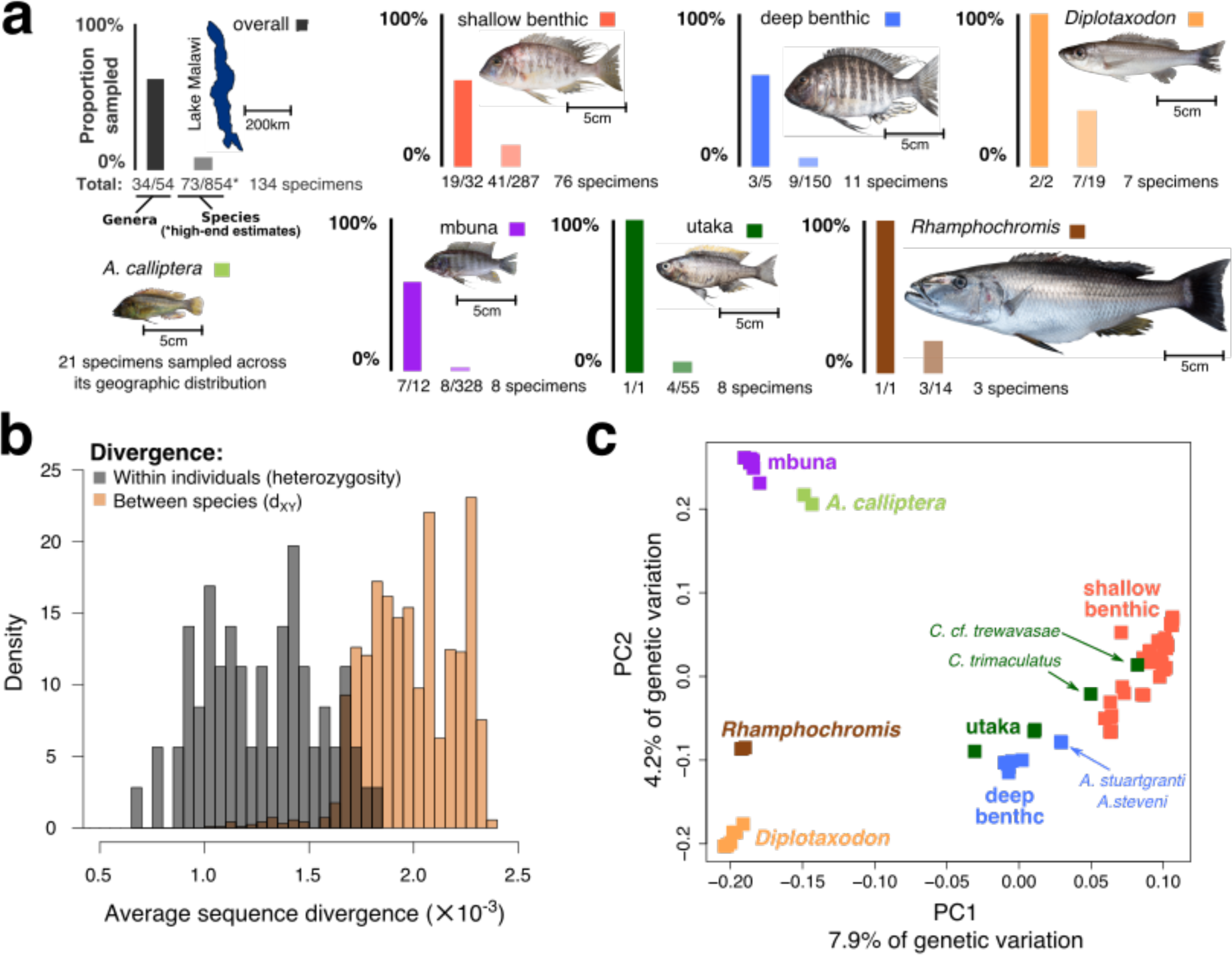
The Lake Malawi haplochromine cichlid radiation. **a,** The sampling coverage of this study: overall and for each of the seven main eco-morphological groups within the radiation. A representative specimen is shown for each group *(Diplotaxodon: D. limnothrissa*; shallow benthic: *Lethrinops albus*; deep benthic: *Lethrinops gossei;* mbuna: *Metriaclima zebra;* utaka: *Copadichromis virginalis*; *Rhamphochromis: R. woodi).* Numbers of species and genera are based on ref. ^22^ **b,** The distributions of genomic sequence diversity within individuals (heterozygosity; *π)* and of divergence between species (d_XY_). **c,** Principal component analysis (PCA) of whole genome variation data.

Despite their extensive phenotypic differentiation, all species within the Lake Malawi haplochromine cichlid radiation are genetically closely related^23,24^. However, genome-wide genetic divergence has never been quantified. To examine the extent of genetic differentiation between species across the radiation we calculated the average pairwise differences between all pairs of sequences from different species (d_XY_). A comparison of d_XY_ against heterozygosity reveals that the two distributions partially overlap (Fig. 1b). Thus, the sequence divergence within a single diploid individual is sometimes higher than the divergence between sequences found in two distinct species. The average d_XY_ is 2.0× 10^−3^ with a range between 1.0 and 2.4×10^−3^ per bp. The maximum d_XY_ is therefore approximately one fifth of the divergence between human and chimpanzee^25−3^. The low ratio of divergence to diversity means that most genetic variation is shared between species. On average both alleles are observed in other species for 82% of heterozygous sites within individuals, consistent with the expected and previously observed high levels of incomplete lineage sorting (ILS) between Lake Malawi species^24^. Supplementary Fig. 2 shows average d_XY_ and F_ST_ values based on comparisons between individuals from the seven eco-morphological groups.

### Low per-generation mutation rate

It has been suggested that the species richness and morphological diversity of teleosts in general and of cichlids in particular might be explained by elevated mutation rates compared to other vertebrates^26,27^. To obtain a direct estimate of the per-generation mutation rate, we reared offspring of three species from three different Lake Malawi groups *(Astatotilapia calliptera, Aulonocara stuartgranti*, and *Lethrinops lethrinus).* We sequenced both parents and one offspring of each to high coverage (40x), applied stringent quality filtering, and counted variants present in each offspring but absent in both its parents (Supplementary Fig. 3 and Supplementary Methods). There was no evidence for significant difference in mutation rates between species. The overall mutation rate (μ) was estimated at 3.5× 10^−9^ (95%CI: 1.6× 10^−9^ to 4.6×10^−9^) per bp per generation, approximately three to four times lower than the rate obtained in similar studies using human trios^28^, although given much shorter mean generation times the per-year rate is still expected to be higher in cichlids than in humans. We note that Recknagel et al.^27^ obtained a much higher mutation rate estimate (6.6× 10^−8^ per bp per generation) in Midas cichlids, but from relatively low depth RADseq data that may have made accurate verification more difficult. We also note that our per generation rate estimate, although low, is still higher than the lowest μ estimate in vertebrates: 2×10^−9^ per bp per generation recently reported for Atlantic herring^29^. By combining our mutation rate with nucleotide diversity (π) values, we estimate the long term effective population sizes (N_e_) to be in the range of approximately 50,000 to 130,000 breeding individuals (with N_e_ = π/4µ). Given previous estimates of the age of the radiation of the order of a million years^12,30,31^ (max estimate 4.63 million years^32^), and the hundreds of species present, this result suggests that alleles at a locus will only rarely coalesce within the time between successive speciation events, consistent with high sharing of heterozygosity and ILS. This is because both the mean and standard deviation in the time to the common ancestor of a pair of alleles are expected to be in the order of 2Ne generations, or hundreds of thousands of years.

### Genome data support for eco-morphological groupings

Principal component analysis (PCA; Fig. 1c) performed on the whole-genome genotype data generally separates the major eco-morphological groups. The most notable exceptions to this are (1) the utaka, for which some species cluster more closely with deep benthics and others with shallow benthics, and (2) two species of the genus *Aulonacara, A. stuartgranti* and *A. steveni*, which are located between the shallow and deep benthic groups. Although these have enlarged lateral line sensory apparatus like many deep benthic species including other *Aulonocara*, they are typically found in shallower water^22^. Another interesting pattern in the PCA plot is the extension of all the utaka and benthic samples towards *Diplotaxodon* (Fig. 1c), a pattern typical for admixed populations (e.g. ref. ^33^). In particular among the benthics, deeper water species are closer to the deep water *Diplotaxodon*, an observation we will return to below in the context of gene flow and shared mechanisms of depth adaptation.

As a further test of the consistency of group assignments, we calculated for each species S all Patterson’s D statistics (ABBA-BABA test; see Supplementary Methods)^34^, of the form D(P1, P2; S, O) where P2 samples were from the same eco-morphological group as S, P1 samples were from a different group, and the outgroup O was fixed as *Neolamprologus brichardi* from Lake Tanganyika. Positive D values reflect excess sharing of derived alleles between S and P2, compared to P1. Therefore, under correct group assignment for S, we would expect all D(P1, P2; S, O) to be positive. The results of this analysis suggest that group assignments were fully supported except for the four species also highlighted in the PCA: the two shallow-living *Aulonocara* species mentioned above are closer to shallow benthics than to deep benthics in 71% and 82% respectively of tests when comparing these alternatives, and *Copadichromis trimaculatus* is closer to shallow benthics than to utaka in 58% of the comparisons involving these two groups. Finally, *Copadichromis cf. trewavasae* clustered with shallow benthics all comparisons; therefore this particular *Copadichromis* species is treated as a member of the shallow benthic group throughout the remainder of this manuscript. Once the three intermediate samples were removed and *C. cf. trewavasae* reassigned, all other species showed 100% consistency with their group assignment.

The overall sub-structuring into groups is also supported by patterns of linkage disequilibrium (LD). Mean LD decays within a few hundred base-pairs across the set of all species, in a few kilobases (kb) for subsets of species from within eco-morphological groups, and extends beyond 10kb within species (Supplementary Fig. 4 and Supplementary Methods).

### Allele sharing reveals that many relationships are not tree-like

The above observations suggest that some species may be genetic intermediates between well-defined groups, consistent with previous studies which have suggested that hybridization and introgression subsequent to initial separation of species may have played a significant role in cichlid radiations, including in Lakes Tanganyika^12,14–16^ and Malawi^18,20^. Where this happens there is no single tree relating the species.

To test more generally for the extent of violation of tree-like species relationships, we calculated Patterson’s D statistic for all possible trios of Lake Malawi species, without assuming any *a priori* knowledge of their relationships. For each trio of species, (A,B,C), we calculated the statistic for all three possible tree topologies with the outgroup again fixed as *N. brichardi* from Lake Tanganyika. The test statistic Dmin is the minimum absolute value of these: D_min_=min(|D(A,B;C,O)|, |D(A,C;B,O)|, |D(C,B;A,O)|). In this case, a significantly positive D_min_ score (see Supplementary Methods) signifies that the sharing of derived alleles between the three species is inconsistent with any tree relating them, even in the presence of incomplete lineage sorting.

The analysis revealed that non-tree-like relationships are pervasive throughout the dataset. Overall, the sharing of derived alleles in 62% of trios (75,616 out of 121,485) is inconsistent with a single species tree relating them (Holm-Bonferroni FWER < 0.01). Our approach is conservative in the sense that the Dmin score for each triplet is considered in isolation and we ignore ‘higher-order’ inconsistencies where D_min_ triplet topologies are inconsistent with each other. We note however that Dmin values are not independent and that a single gene flow event between ancestral lineages can affect multiple contemporary species and thus affect more triplets than a more recent gene flow event would. Strong tree violations are seen within all the major groups and also between groups (Fig. 2a), suggesting reticulate evolution at multiple levels, and that an accurate description of the evolutionary relationships of Lake Malawi cichlids requires some form of network, rather than a phylogenetic tree.

**Fig. 2:**
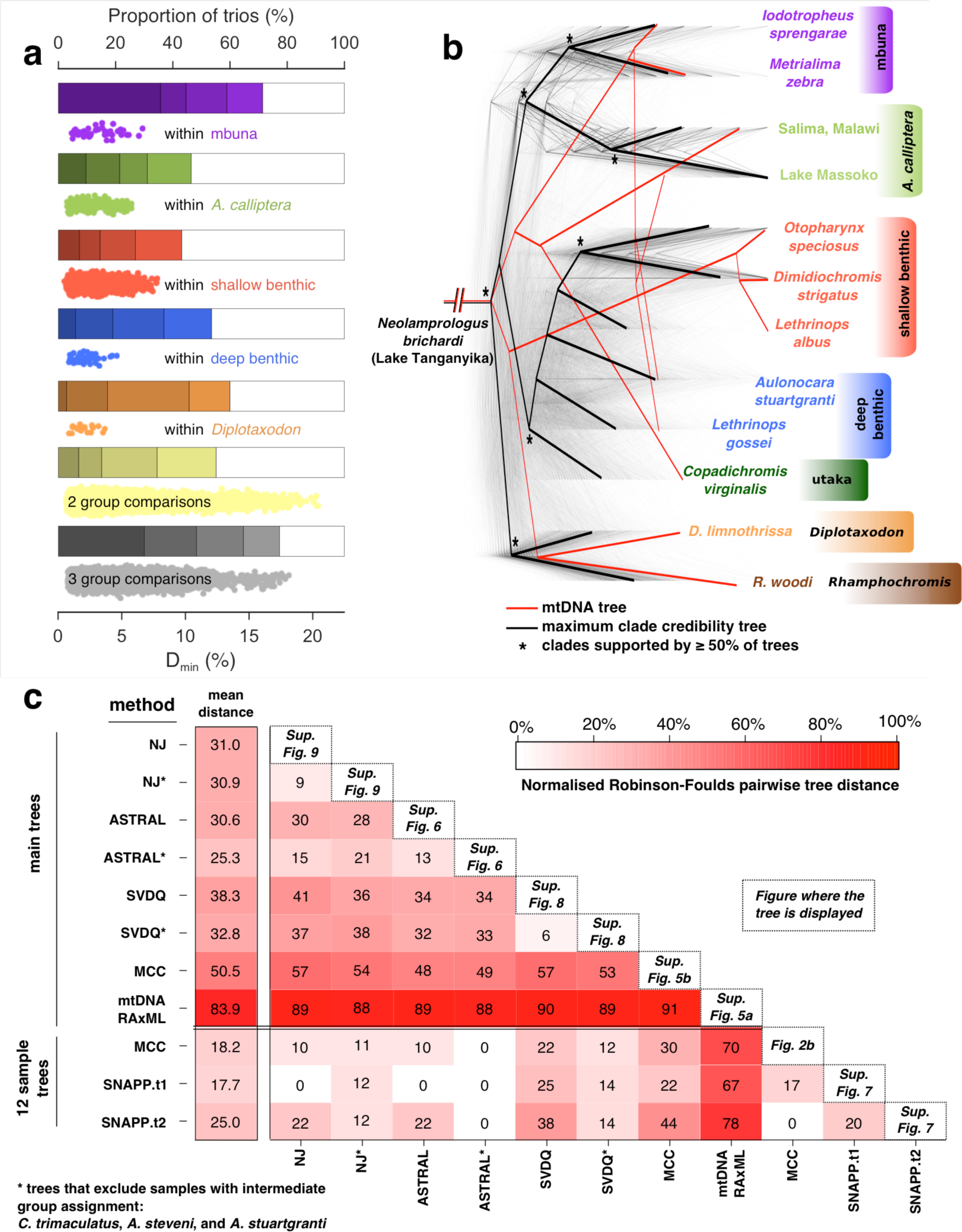
Excess allele sharing and patterns of species relatedness. **a**, Derived allele sharing reveals non-tree-like relationships among trios of Lake Malawi cichlid species. The bars show the proportion of significantly elevated D_min_ scores (defined as the smallest |D| statistics obtainable for each trio of species with outgroup *Neolamprologus brichardi).* Shading corresponds to FWER q values of (from light to dark) 10^−2^, 10^−4^, 10^−8^, 10^−14^. The scatterplots show the Dmin scores that were significant at FWER<0.01. Results are shown separately for within-group comparisons, where all three species in the trio are from the same group, and for cases where the species come from two or three different groups. *Rhamphochromis* and utaka within-group comparisons are not shown due to the low number of data points. **b**, A set of 2543 Maximum Likelihood (ML) phylogenetic trees for non-overlapping regions along the genome. Branch lengths were scaled for visualization so that the total height of each tree is the same. The local trees were built with 71 species and then subsampled for display to 12 individuals representing the eco-morphological groups. The maximum clade credibility tree shown here was built from the region phylogenies with 12 individuals. An ML mitochondrial phylogeny is shown for comparison. **c**, A summary of phylogenetic trees produced in this study and the normalised Robinson-Foulds distances between them, reflecting the topological distance between pairs of trees on the scale from zero to 100%. The mtDNA tree is a clear outlier with the highest topological distance to all other trees (mean=83.9%). The least controversial 12 sample tree is SNAPP.t1, with an average distance to other trees of 17.7%, while ASTRAL* is the least controversial among the ‘main trees’ (mean distance of 25.3%). To compare trees with differing sets of taxa, the trees were downsampled so that only matching taxa were present for the comparisons. The position of the outgroup, i.e. the root of the radiation, was always considered in all tree comparisons. See Supplementary Methods for details.

### Phylogenetic framework relationships

Despite no tree giving a complete and accurate picture of the relationship between species, standard phylogenetic approaches are useful to provide a framework for discussion. To obtain an initial picture we divided the genome into 2543 non-overlapping windows, each comprising 8000 SNPs (average size: 274kb), and constructed a Maximum Likelihood (ML) phylogeny separately for the full sequences within each window, obtaining trees with 2542 different topologies (see Supplementary Methods). We also calculated the maximum clade credibility (MCC) summary tree^35^ and an ML phylogeny based on the full mtDNA genome (Fig. 1c and Supplementary Fig. 5).

We next applied a range of further phylogenomic methods which are known to be robust to incomplete lineage sorting. These included three multispecies coalescent methods^36,37^: the Bayesian SNAPP^38^ (with a subset of 48,922 unlinked SNPs in 12 individuals representing the eco-morphological groups), the algebraic method SVDquartets^39,40^, which allows for site-specific rate variation and is robust gene-flow between sister taxa^41^, and the summary method ASTRAL^42,43^, using the 2543 local ML trees that were described above as input. We also built whole genome Neighbour-Joining (NJ) tree using the Dasarathy et al. algorithm, which has been shown to be a statistically consistent and accurate species tree estimator under ILS^44,45^. The above methods have also been applied to datasets where the individuals that are genetically intermediate between eco-morphological groups *(C. trimaculatus, A. stuartgranti*, and *A. steveni)* have been removed, thus likely reducing the extent of violation of the multispecies coalescent model.

Despite extensive variation among the 2543 individual ML trees (at least in part attributable to ILS), and, to a lesser extent, variation between the different genome-wide phylogenetic methods, there is some general consensus (Fig. 2c and Supplementary Figs. 5-9). Except for the three previously identified intermediate species, individuals from within each of the previously identified eco-morphological groups cluster together in all the whole genome phylogenies, forming well supported reciprocally monophyletic groups. The pelagic *Diplotaxodon* and *Rhamphochromis* together form a sister group to the rest of the radiation, except in the all-sample MCC and SVDquartets phylogenies. Perhaps surprisingly, all the methods place the widely-distributed lake/river-dwelling *A. calliptera* as the sister taxon to the specialized rocky-shore mbuna group in a position that is nested within the radiation. On a finer scale, many similarities between the resulting phylogenies reflect features of previous taxonomic assignment, but some currently-recognized genera appear in all the trees as clearly polyphyletic, including *Placidochromis, Lethrinops*, and *Mylochromis.*

It is also clear that the mtDNA phylogeny is an outlier, being substantially different from all the whole-genome phylogenies and also from the majority of the local ML trees (Fig. 2b,c and Supplementary Figs. 5 and 10). Discordances between mtDNA and nuclear phylogenies in Lake Malawi have been reported previously^18,20^ and interpreted as a signature of past hybridization events. However, large discrepancies between mitochondrial and nuclear phylogenies have been shown in many other systems, reflecting both that mtDNA as a single locus is not expected to reflect the consensus under ILS, and that it often does not evolve neutrally (e.g. refs. ^46–48^). In particular, the high incidence of mitochondrial selection underlines the importance of evaluating the Lake Malawi radiation from a genome-wide perspective rather than drawing conclusions regarding species relationships based on mtDNA signals alone. Indeed, as we discuss below, some of the previously suggested hybridization events based on comparisons between nuclear and mitochondrial markers are not reflected in the whole genome data.

### Strong signals of introgression

Next we applied a variety of methods to identify the species and groups whose relationships violate the framework trees described in the previous section. First, we contrasted the pairwise genetic distances used to produce the raw NJ tree against the distances between samples along the tree branches, calculating the residuals (Supplementary Fig. 11). If the tree were able to perfectly capture all the genetic relationships in our sample, we would expect the residuals to be zero. However, we found a large number of differences, with some standout cases.

The patterns of differences between the true genetic distance and the NJ tree distance affect both groups of species and individual species. Among the strongest signals on individual species, in addition to the previously discussed *C. trimaculatus*, we can see that 1) *Placidochromis* cf. *longimanus* is genetically closer to the deep benthic clade and to a subset of the shallow benthic (mainly *Lethrinops* species) than the tree suggests; and 2) our sample of *Otopharynx tetrastigma* is much closer to *Astatotilapia calliptera* from Lake Kingiri (and to a lesser degree to other *A. calliptera*) than would be expected from the tree. The *O. tetrastigma* specimen comes from Lake Ilamba, a satellite crater lake of Lake Malawi that also harbours a population of *A. calliptera* and is geographically close (3.2 km) to Lake Kingiri.

Second, the sharing of long haplotypes between otherwise distantly related species is an indication of recent admixture or introgression. To investigate this type of gene flow signature, we used the chromopainter software package^49^ and calculated a ‘coancestry matrix’ of all species - a summary of nearest neighbour (therefore recent) haplotype relationships in the dataset (see Supplementary Methods). We found that the Lake Ilamba *O. tetrastigma* and Lake Kingiri *A. calliptera* stand out again in this analysis as showing a strong signature of recent gene flow between individual species from distinct eco-morphological groups (Supplementary Fig. 12). The other tree-violation signatures described above are also visible on the haplotype sharing level but are less pronounced, consistent with being older events perhaps involving the common ancestors of multiple present-day species. However, the chromopainter analysis reveals numerous other examples of excess co-ancestry between species that do not cluster immediately together (e.g. the utaka *C. virginalis* with *Diplotaxodon*; more highlighted in Supplementary Fig. 12). Furthermore, the clustering based on recent co-ancestry is different from any tree generated using phylogenetic methods; in particular a number of shallow benthics including *P*. cf. *longimanus* cluster next to the deep benthics.

Third, we used the *f*_4_ admixture ratio^34,50,51^ (*f* statistic in the following), a statistic closely related to Patterson’s D^34^, computing *f*(A,B;C,O) for all trios of species that fit the relationships ((A, B), C) in the ASTRAL* tree from Supplementary Figure 6, with the outgroup fixed as *N. brichardi.* The ASTRAL* tree has the lowest mean topological distance to all the other trees, and excludes the three species with intermediate group assignment, a choice made here because we were interested in identifying additional signals beyond the admixed status of *A. stuartgranti, A. steveni*, and *C. trimaculatus.*

When elevated due to introgression the *f* statistic is expected to be linear in relation to the proportion of introgressed material. Out of 164,320 computed *f* statistics 97,889 were significant at FWER < 0.001. However, as was the case for the Patterson’s D statistics presented above, a single gene flow event can lead to multiple significant *f* statistics, so the values calculated for different combinations of ((A, B), C) groups are not independent as soon as they share internal or external branches. Therefore, we sought to obtain branch-specific estimates of excess allele sharing that would be less correlated. Building on the logic employed to understand correlated gene flow signals in ref. ^52^, we developed “f-branch” or *f*_*b*_*(C):* a summary of *f* scores that captures excess allele sharing between a clade *C* and a branch *b* compared to the sister branch of *b* (see Supplementary Methods). Therefore, the *f*_*b*_*(C)* scores are specific for a branch *b* (on the y-axis in Fig. 3), but a single introgression event can still lead to significant *f*_*b*_(*C*) for multiple related C. In simulations *f-*branch is more robust than treemix^53^ (Supplementary Note).

**Fig. 3:**
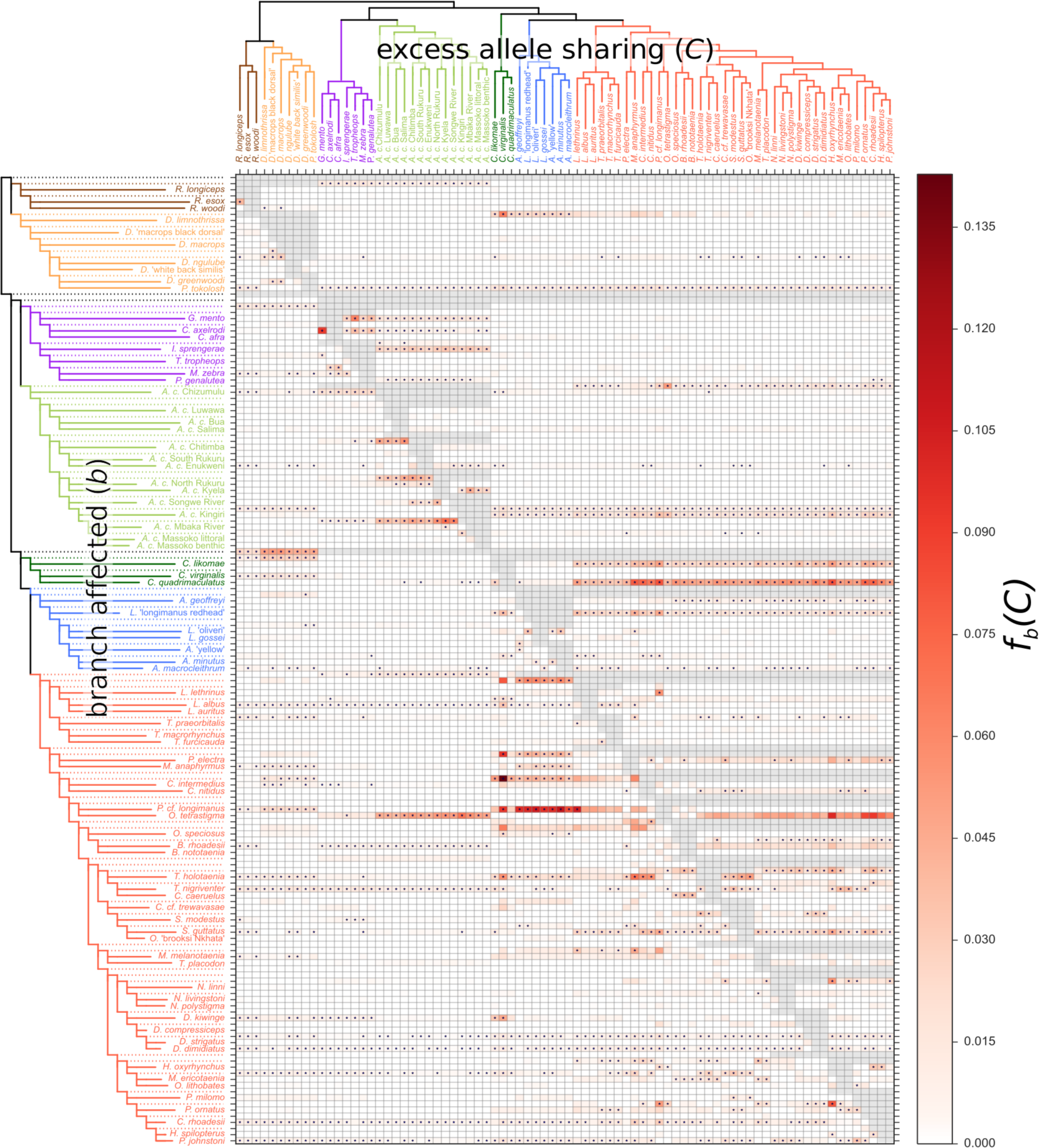
Identifying tree violating branches and possible gene flow events. The branch-specific statistic *f*_*b*_*(C)* identifies excess sharing of derived alleles between the branch of the tree on the y-axis and the species C on the x-axis (see Supplementary Note). The ASTRAL* tree was used as a basis for the branch statistic and grey data points in the matrix correspond to tests that are not consistent with the phylogeny. Colours correspond to eco-morphological groups as in Fig. 1. The * sign denotes block jack-knifing significance at |Z|>3.17 (Holm-Bonferroni FWER<0.001).

The *f*-branch summary reduces the number of *f* statistics more than tenfold, to 11,158. Of these, 1,421 are still significantly elevated (at FWER<0.001; Supplementary Fig. 13), while 92 of the 158 branches in the phylogeny show significant excess allele sharing with at least one other species *C* (Fig. 3). Not only are there many significant *f*_*b*_*(C)* scores, they also are unusually large: 238 out of the 1,421 significant scores (2.1% of the total 11,158) are larger than 3%, more than inferred for human-Neanderthal introgression^34^. The *f* statistic tests are robust to the occurrence of incomplete lineage sorting, in the sense that ILS alone cannot generate a significant test result^50^. We note, however, that pronounced population structure within ancestral species, coupled with rapid succession of speciation events, can also substantially violate the assumptions of a strictly bifurcating species tree and lead to significantly elevated *f* scores^50,54^. This needs to be taken into account when interpreting non-treelike relationships, for example among major groups early in the radiation. However, in cases of excess allele sharing between ‘distant’ lineages that are separated by multiple speciation events, ancestral population structure would have needed to segregate through these speciation events without affecting sister lineages, a scenario that is not credible in general. Therefore, we suggest that there is strong evidence for multiple cross-species gene flow events.

The overall strongest signal of excess allele sharing *(f*_*b*_*(C)* = 14.2%) is between the ancestral lineage of the two sampled *Ctenopharynx* species from the shallow benthic group and the utaka *Copadichromis virginalis* (Fig. 3). Notably, *Ctenopharynx* species, particularly *C. intermedius* and *C. pictus* have very large numbers of long slender gill rakers, a feature shared with *Copadichromis* species, and believed to be related to a diet of small invertebrates^55^. Several other benthic lineages also share excess alleles with *C. virginalis* at a lower level. Next, the significantly elevated *f*_*b*_*(C)* scores between the shallow and the deep benthic lineages suggest genetic exchange between these two groups go beyond the clearly admixed shallow-living *Aulonacara* (not included in this analysis). The f-branch signals between *O. tetrastigma* and *A. calliptera* Kingiri are observed in both directions – *A. calliptera* Kingiri with shallow benthics (and most strongly *O. tetrastigma)* and *O. tetrastigma* with *A. calliptera* (most strongly *A. c.* Kingiri), suggesting bi-directional introgression.

At the level of the major eco-morphological groups, the strongest signal is that the ancestral lineage of benthics and utaka shares excess derived alleles with *Diplotaxodon* and, to a lesser degree, *Rhamphochromis*, as previously suggested by the PCA plot (Fig 1c). Furthermore, there is evidence for additional ancestry from the pelagic groups in utaka, which could be explained either by an additional, more recent, gene flow event or by differential fixation of introgressed material, possibly due to selection. Reciprocally, *Diplotaxodon* shares excess derived alleles (relative to *Rhamphochromis)* with utaka and deep benthics, as does *Rhamphochromis* with mbuna and *A. calliptera.* Furthermore, mbuna show excess allele sharing (relative to *A. calliptera)* with *Diplotaxodon* and *Rhamphochromis.* (Fig. 3) On the other hand, while ref. ^18^ suggested gene flow between the deep benthic and mbuna groups on the basis of mtDNA phylogeny, our genome-wide analysis did not find any signal of substantial genetic exchange between these groups.

Overall, the D statistics, the NJ tree residuals, the haplotype sharing patterns, and the many elevated *fb(C)* scores in Fig. 3 reveal extensive violations of the bifurcating species tree model both across and within major groups. Considering the magnitude and patterns of these violations, we conclude that no single species tree can adequately capture the evolutionary relationships within the species flock and that there were multiple gene flow events at different times during the evolutionary history of the adaptive radiation, potentially with an additional contribution of ancestral population structure.

### Origins of the radiation

*Astatotilapia calliptera*, although showing strong preference for shallow weedy habitats, is abundant and widespread in both flowing and still waters, and is thus considered a generalist. It has often been referred to as the ‘prototype’ for the closely-related endemic genera of Lake Malawi cichlids^22^. Therefore, discussions concerning the origin of the Malawi radiation often rely on ascertaining the relationship of this species to the rest of the Malawi radiation^56^. Previous phylogenetic analyses, using mtDNA and small numbers of nuclear markers, showed inconsistencies with respect to *A. calliptera* (compare refs. ^20^ and ^18^). On the other hand, our whole genome data indicate a clear and consistent position of all the *A. calliptera* individuals from the Lake Malawi catchment as members of a sister group to the mbuna, consistent with the nuclear DNA phylogeny in ref. ^18^, although that study included only a single *A. calliptera* specimen.

To explore the origins of the Lake Malawi radiation in greater detail, we obtained whole genome sequences from 19 individuals from seven *Astatotilapia* species not found in Lake Malawi (Supplementary Table 2) and generated new variant calls (Supplementary Methods). In addition, we sequenced five more *A. calliptera* specimens from Indian Ocean catchments, thus covering most of the geographical distribution of the species. We constructed NJ trees based on genetic distances and found that even with these additional data all the *A. calliptera* (including samples from outside the Lake Malawi catchment) continue to cluster as a single group nested at the same place within the radiation, whereas the other *Astatotilapia* species clearly branched off well before the lake radiation (Fig. 4a,b,c).

Joyce et al.^20^ reported that the mtDNA haplogroup of *A. calliptera* from the Indian Ocean catchment clustered with mbuna (as we see for mtDNA tree in Supplementary Fig. 14) and concluded that the phylogenetic discordances between mtDNA and nuclear markers can be explained by gene-flow. They suggested that there had been repeated colonization of Lake Malawi by independent *Astatotilapia* lineages with different mitochondrial haplogroups, the first founding the entire species flock, and the second, with the mtDNA haplogroup common in the Indian Ocean catchment, introgressing into the Malawi radiation and contributing strongly to the mbuna group. This hypothesis predicts that among the *A. calliptera*, the Indian Ocean catchment individuals should be closer to mbuna than the individuals sampled within the Malawi catchment. However, using the *f* statistics, we found a strong signal in the opposite direction across the nuclear genome *f*= 30% for Malawi catchment *A. calliptera* being closer to mbuna; Fig. 4d). Therefore, the Joyce et al. hypothesis that the mbuna, the most species rich group within the radiation, may be a hybrid lineage formed from independent invasions is not supported by genome-wide data.

It has been repeatedly suggested that *A. calliptera* may be the direct descendant of the riverine-generalist lineage that seeded the Lake Malawi radiation [e.g. refs. ^7,55–57^]. Our interpretation of this argument is that *A. calliptera* itself is not the ancestor of the radiation, but it is likely that the ancestor was a riverine-generalist that was ecologically and phenotypically similar or even equivalent to it. This hypothesis is lent further support by our geometric morphometric analysis. Using 17 homologous body shape landmarks (Supplementary Methods) we established that, despite the relatively large genetic divergence, *A. calliptera* is nested within the morphospace of the other more distantly related but ecologically similar *Astatotilapia* species (Fig. 4a,e), and these together have a central position within the morphological space of the Lake Malawi radiation (Fig. 4e and Supplementary Fig. 15).

**Fig. 4:**
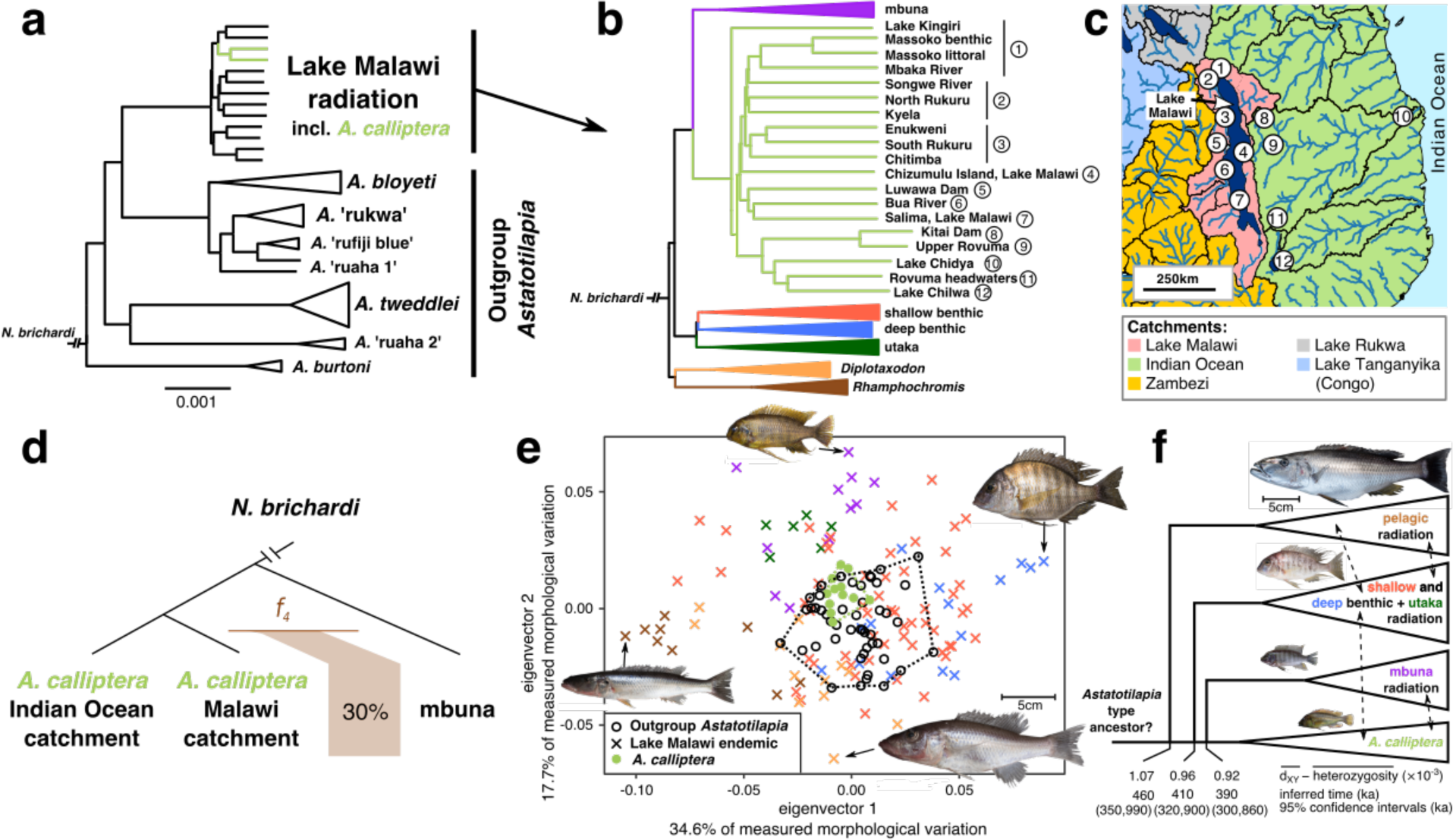
Origins of the radiation and the role of *A. calliptera.* **a**, An NJ phylogeny showing the Lake Malawi radiation in the context of other East African *Astatotilapia* taxa. **b**, A Lake Malawi NJ phylogeny with expanded view of *A. calliptera*, with all other groups collapsed. **c**, Approximate *A. calliptera* sampling locations shown on a map of the broader Lake Malawi region. Black lines correspond to present day level 3 catchment boundaries from the US Geological Survey’s HYDRO1k dataset. **d**, Strong *f*_*4*_ admixture ratio signal showing that Malawi catchment *A. calliptera* are closer to mbuna than their Indian Ocean catchment counterparts. **e**, PCA of body shape variation of Lake Malawi endemics, *A. calliptera* and other *Astatotilapia* taxa, obtained from geometric morphometric analysis. **f**, A phylogeny with the same topology as in panel (b) but displayed with a straight line between the ancestor and *A. calliptera.* For each branch off this lineage, we show mean sequence divergence (d_XY_) minus mean heterozygosity, translation of this value into a mean time estimate, and 95% CI for the time estimate reflecting the statistical uncertainty in mutation rate. Dashed lines with arrows indicate likely instances of gene flow between major groups; their absolute timing (position along the x axis) is arbitrary

Therefore, we propose here a model which reconciles the nested phylogenetic position of *A. calliptera* in all of the whole genome phylogenies with its generalist riverine phenotype. We suggest that the Lake Malawi species flock consists of three separate radiations splitting off from the lineage leading to *A. calliptera*. The relationships between the major groups supported by the ASTRAL, SNAPP and NJ methods suggest that the pelagic radiation was seeded first, then the benthic + utaka, and finally the rock-dwelling mbuna, all in a relatively quick succession, followed by subsequent gene flow as described above (Fig. 4f; note the pelagic vs. utaka + benthic branching order is swapped in SVDquartets tree as shown in Supplementary Fig. 8b). Applying our per-generation mutation rate to observed genomic divergences we obtained mean separation time estimates between these lineages between 460 thousand years ago (ka) [95%CI: (350ka to 990ka)] and 390ky [95%CI: (300ka to 860ka)] (Fig. 4f), assuming three years per generation as in ref. ^58^. The point estimates all fall within the second most recent prolonged deep lake phase as inferred from the Lake Malawi paleoecological record^30^ while the upper ends of the confidence intervals cover the third deep lake phase. We also note that split times estimated from sequence divergence are likely to be reduced by subsequent gene-flow, leading to underestimates. Therefore we conclude that the data are consistent with the previous reports based on fossil time calibration which put the origin of the Lake Malawi radiation at 700-800ka^12^.

Focusing on the *A. calliptera* individuals, we found they cluster by geography (Fig. 4b,c), except for the specimen from crater Lake Kingiri, whose position in the tree is likely a result of the admixture signals shown in Fig. 3. Indeed, the Kingiri individual clusters according to geography with the specimens from the nearby crater Lake Massoko and Mbaka River if a NJ tree is built with *A. calliptera* samples only (Supplementary Fig. 16). Applying the same logic, we tested whether the position of the *A. calliptera* group in the NJ tree changes when the tree is built without mbuna (as would be expected if the *A. calliptera* position were affected by hybridization with mbuna). However, we found that the position of *A. calliptera* is not affected by the removal of mbuna (Supplementary Fig. 17), suggesting that the nested position is not due to later hybridization. The *f* statistic analysis in Fig. 3 further supports this claim, because the signals involving the whole mbuna or *A. calliptera* groups are modest and do not suggest erroneous placement of these whole groups in all phylogenetic analyses.

Furthermore, the nested position of *A. calliptera* is also supported by the vast majority of the genome. We specifically searched for the basal branch in a set of 2638 local ML phylogenies for non-overlapping genomic windows and found results that are consistent with the whole genome ASTRAL, SNAPP and NJ trees: the most common basal branches are the pelagic groups *Rhamphochromis* and *Diplotaxodon* (in 42.12% of the genomic windows). In comparison, *A. calliptera* (including all of the Indian Ocean catchment samples) were found to be basal only in 5.99% of the windows (Supplementary Fig. 18).

Finally, we note that all the *A. calliptera* have a relatively recent common ancestor, with divergence at ~75% of the most distinct species in the Malawi radiation and corresponding to 340ka [95%CI: (260ka, 740ka)], suggesting that the Lake Malawi population has been a reservoir that has repopulated the river systems and more transient lakes following dry-wet transitions in East African hydroclimate^30,59^. Our results do not fully resolve whether the lineage leading from the common ancestor to *A. calliptera* retained its riverine generalist phenotype throughout or whether a lacustrine species evolved at some point (e.g. the common ancestor of *A. calliptera* and mbuna) and later de-specialized again to recolonize the rivers. However, while it is a possibility, we suggest that is it not likely that the many strong phenotypic affinities of *A. calliptera* to the basal *Astatotilapia* [see refs. ^60,61^; Fig. 4e], would be reinvented from a lacustrine species. After all, *A. calliptera* is widespread and abundant in its preferred shallow weedy habitat throughout present-day Lake Malawi, despite the presence of hundreds of closely-related endemic lacustrine specialist species.

### Signatures and consequences of selection on coding sequences

To gain insight into the functional basis of diversification and adaptation in Lake Malawi cichlids, we next turned our attention to protein coding genes. We compared the between-species levels of non-synonymous variation 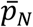 to synonymous variation 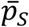 in 20,664 genes and calculated the difference between these two values (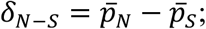 Supplementary Methods). Overall, coding sequence exhibits signatures of purifying selection: the average between-species 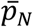 was 54% lower than in a random matching set of non-coding regions. Interestingly, the average between-species synonymous variation 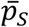 in genes was slightly but significantly higher than in non-coding control regions (13% lower mean; *p <* 2.2×10^−16^, one tailed Mann-Whitney test). One possible explanation of this observation would be if intergenic regions were homogenized by gene-flow, whereas protein coding genes were more resistant to this.

To control for statistical effects stemming from variation in gene length and sequence composition we normalized the *δ*_*N-S*_ values per gene by taking into account the variance in *p*_*N*_ — *p*_*S*_ across all pairwise sequence comparisons for each gene, deriving the non-synonymous excess score (∆_*N-S*_.; Supplementary Methods). We focus below on the top 5% of the ∆_*N-S*_ distribution (∆_*N-S*_ > 40.2, 1034 candidate genes); these high values at the upper tail of the distribution are substantially overrepresented in the actual data when compared to a null model based on random sampling of codons (Fig. 5a; Supplementary Methods). To obtain a highly positive ∆_*N-S*_ genes need to have been under positive selection at multiple non-synonymous sites, either recently repeatedly within multiple species or ancestrally. Therefore, it reveals only a limited subset of positive selection events that occurred in the history of the radiation (e.g. a single selection event on a single amino acid would not be detected).

This candidate gene set is highly enriched for genes for which no homologs were found in any of medaka, stickleback, *Tetraodon* or zebrafish (other teleosts) when examined in ref. ^11^ (606 out of 4,190 without vs. 428 out of 16,472 with homology assignment; *χ*^2^ test *p* < 2.2* 10^−16^). Genes without homologs tend to be short (median coding length is 432bp) and some of the signal may be explained by a component of gene prediction errors. However a comparison of short genes (≤450bp) without homologs to a set of random noncoding sequences (Supplementary Fig. 19) showed significant differences (*p* < 2.2× 10^−16^, Mann-Whitney test), with both a substantial component of genes with low 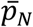, reflecting genes under purifying selection, and also an excess of genes with high 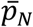 (Supplementary Fig. 20).

Cichlids have an unexpectedly large number of gene duplicates and it has been suggested that this phenomenon has contributed to their extensive adaptive radiations^3,11^. To investigate the extent of divergent selection on gene duplicates, we examined how the non-synonymous excess scores are related to gene copy numbers in the reference genomes. Focusing on homologous genes annotated both in the Malawi reference (*M. zebra*) and in the zebrafish genome, we found that the highest proportion of candidate genes was among genes with two or more copies in both genomes (N - N). The relative enrichment in this category is both substantial and highly significant (Fig. 5b). On the other hand, the increase in proportion of candidate genes in the N - 1 category (multiple copies in the *M. zebra* genome but only one copy in zebrafish) is of a much lesser magnitude and is not significant (*Χ*^2^ test *p* < 0.18), suggesting that selection is occurring more often within ancient multi-copy gene families, rather than on genes with cichlid-specific duplications.

**Fig. 5:**
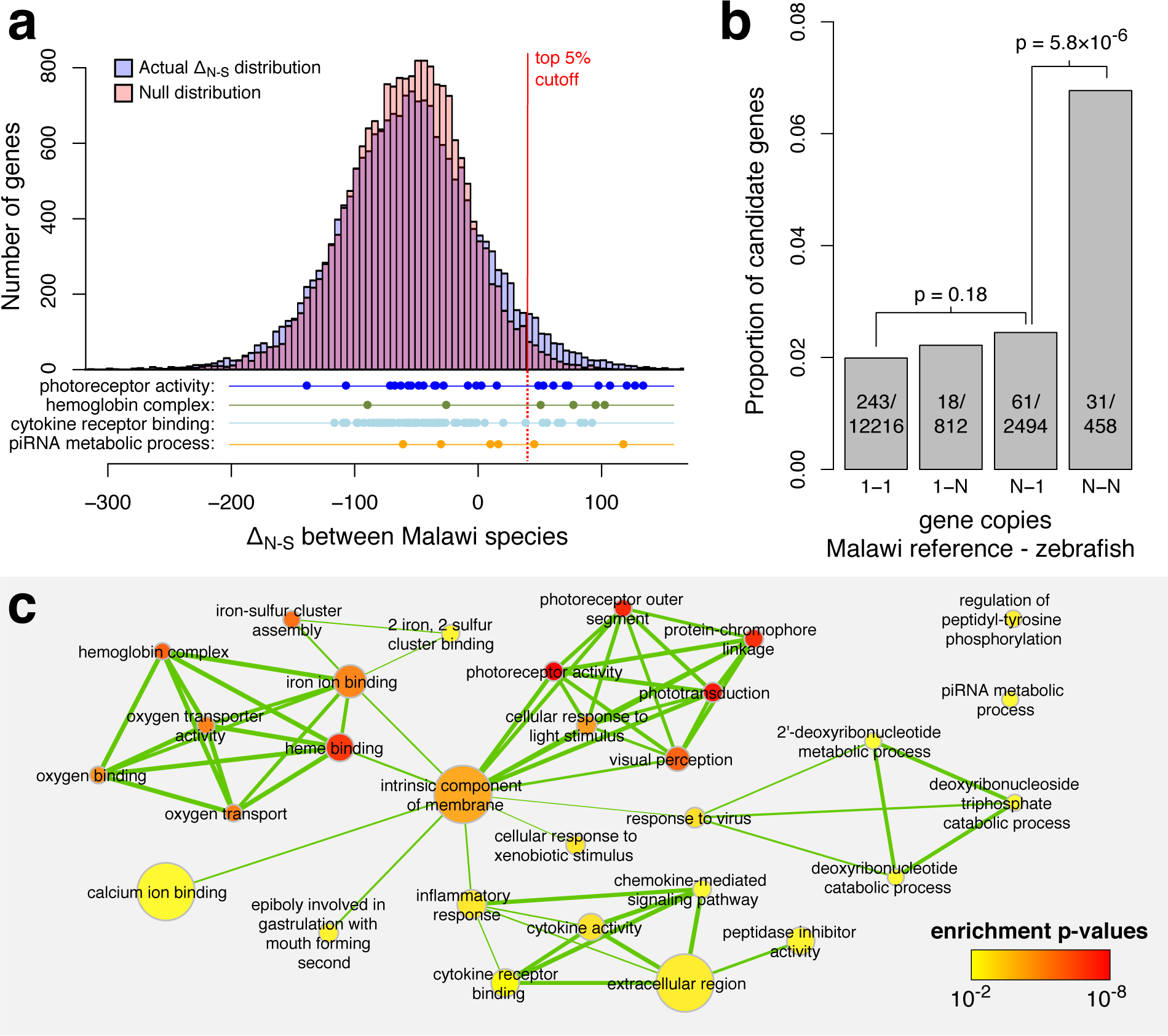
Gene selection scores, copy numbers, and ontology enrichment. **a**, The distribution of the non-synonymous variation excess scores (**∆_*N-S*_**) highlighting the top 5% cutoff, compared against a null model. The null was derived by calculating the statistic on randomly sampled combinations of codons. We also show the distributions of genes in selected Gene Ontology (GO) categories which are overrepresented in the top 5%. **b**, The relationship between the probability of **∆_*N-S*_** being positive and in the top 5% and the relative copy numbers of genes in the Lake Malawi reference (*M. zebra)* and zebrafish. The p-values are based on *Χ*^2^ tests of independence. Genes existing in two or more copies in both zebrafish and Malawi cichlids are disproportionately represented among candidate selected genes. c, An enrichment map for significantly enriched GO terms (cutoff at p ≤ 0.01). The level of overlap between GO enriched terms is indicated by the thickness of the edge between them. The color of each node indicates the p-value for the term and the size of the node is proportional to the number of genes annotated with that GO category.

Next we used Gene Ontology (GO) annotation of zebrafish homologs to test whether candidate genes are enriched for particular functional categories (Supplementary Methods). We found significant enrichment for 30 GO terms (range: 1.6×10^−8^ < p < 0.01, weigh algorithm^62,63^; Supplementary Table 3), 10 in the Molecular Function (MF), 4 in the Cellular Component (CC), and 16 in Biological Process (BP) category. Combining the results from all three GO categories in a network connecting terms with high overlap (i.e. sharing many genes) revealed clear clusters of enriched terms related to (i) haemoglobin function and oxygen transport; (ii) phototransduction and visual perception; and (iii) the immune system, especially inflammatory response and cytokine activity (Fig. 5c). It has been previously suggested that evolution of genes in these functional categories has contributed to cichlid radiations (as discussed below); it is nevertheless interesting to see that these categories stand out in an unbiased analysis of 20,000+ genes.

### Shared mechanisms of depth adaptation

To gain insight into the distribution of adaptive alleles across the radiation, we examined the haplotype relationships for amino acid sequences of candidate genes, focusing on the genes in significantly enriched GO categories, representing potentially complex haplotype genealogy networks by maximum likelihood trees. Many of these haplotype trees in the ‘visual perception’ category have common features that are unusual in the broader dataset: the haplotypes from the deep benthic group and the deep-water pelagic *Diplotaxodon* tend to be disproportionally diverse when compared with the rest of the radiation, and tend to group together despite these two groups being distant in the whole-genome phylogenetic reconstructions. *Diplotaxodon* and deep benthic individuals formed a monophyletic group in only two out of 2638 local ML phylogenies for non-overlapping genomic windows.

Sharply decreasing levels of dissolved oxygen and low light intensities with narrow short wavelength spectra are the hallmarks of the habitats at below ~50 meters to which the deep benthic and pelagic *Diplotaxodon* groups have both adapted, either convergently or in parallel^64^. Signatures of selection on similar haplotypes in the same genes involved in vision and in oxygen transport would therefore point to shared molecular mechanisms underlying this ecological parallelism.

**Fig. 6:**
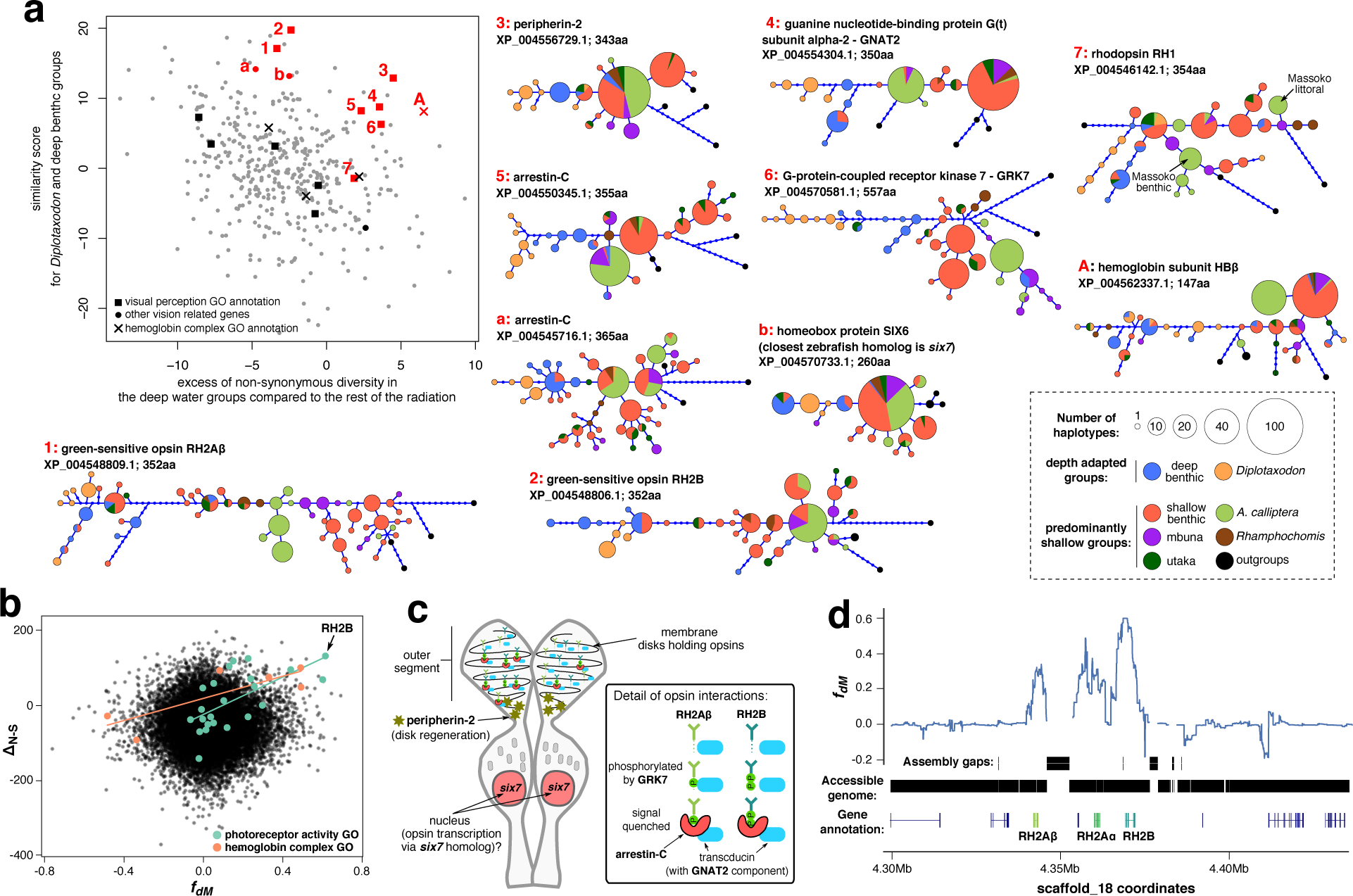
Shared selection between the deep water adapted groups *Diplotaxodon* and deep benthic. **a**, The scatterplot shows the distribution of genes with high ∆_*N-S*_ scores (candidates for positive selection) along axes reflecting shared selection signatures. Only genes with zebrafish homologs are shown. Amino acid haplotype trees, shown for genes as indicated by the red symbols and numbers, indicate that *Diplotaxodon* and deep benthic species are often divergent from other taxa, but similar to each other. Outgroups include *Oreochromis niloticus, Neolamprologus brichardi, Astatotilapia burtoni*, and *Pundamilia nyererei.* **b**, Selection scores plotted against *fdM* (mbuna, deep benthic, *Diplotaxodon, N. brichardi)*, a measure of excess allele sharing between deep benthic and *Diplotaxodon.* Overall there is no correlation between ∆_*N-S*_ and *f*_dM_. However, the strong correlation between ∆_*N-S*_ and *f*_*dM*_ in the highlighted GO categories suggests that positively selected alleles in those categories tend to be subject to introgression between *Diplotaxodon* and the deep benthic group. **c**, A schematic drawing of a double cone photoreceptor expressing the green sensitive opsins and illustrating the functions of other genes with signatures of shared selection. **d**, *f*_*dM*_ calculated in sliding windows of 100 SNPs around the green opsin cluster, revealing that excess allele sharing between deep benthic and *Diplotaxodon* extends far beyond the coding sequences.

To obtain a quantitative measure of shared molecular mechanisms, we calculated for each gene a similarity score for deep benthic and *Diplotaxodon* amino acid sequences and also compared the amounts of non-coding variation in these depth-adapted groups against the rest of the radiation (Supplementary Methods). Both measures are elevated for candidate genes in the ‘visual perception’ category (Fig. 6a; p=0.007 for similarity, p=0.08 for shared diversity, and p=0.003 when similarity and diversity scores are added; all p-values based on Mann-Whitney test). The measures are also elevated for the ‘haemoglobin complex’ category, although due to the small number of genes the differences are not statistically significant in this case. Furthermore, the level of excess allele sharing between *Diplotaxodon* and deep benthic (measured by the local *f* statistic *f*_d*M*_^51,58^) is strongly correlated with the ∆_N-S_ selection score for genes annotated with photoreceptor activity and haemoglobin complex GO terms (*ρ*_*s*_ = 0.63 and 0.81, *p* = 0.001 and *p* = 0.051, respectively, Fig. 6b).

Vision genes with high similarity and diversity scores for the deep benthic and *Diplotaxodon* groups include three opsin genes: the green sensitive opsins RH2Aβ, RH2B, and rhodopsin (Fig. 6a and Supplementary Fig. 21a). The specific residues that distinguish the deep adapted groups from the rest of the radiation differ between the two RH2 copies, with only one shared mutation out of a possible fourteen (Supplementary Fig. 21b). RH2Aβ and RH2B are located within 40kb from each other on the same chromosome (Fig. 6c); a third paralog, RH2Aα, is located between them, but it has very little coding diversity specific to deep benthic and *Diplotaxodon* (Supplementary Fig. 22). This finding is consistent with previous reports suggesting functional divergence between RH2Aα and RH2Aβ following the duplication of RH2A early in the cichlid lineage^65,66^. A similar, albeit weaker signature of shared depth-related selection is apparent in rhodopsin, which is known to play a role in deep-water adaptation in cichlids^67^. Previously, we discussed the role of coding variants in rhodopsin in the early stages of speciation of *A. calliptera* in the crater Lake Massoko^58^. The haplotype tree presented here for the broader radiation strongly suggests that the Massoko alleles did not originate by mutation in that lake but were selected out of ancestral variation (Fig. 6a).

The long wavelength, red-sensitive opsin (LWS) has been shown to play a role in speciation along a depth gradient in Lake Victoria^68^. While it is not particularly diverse in *Diplotaxodon* and deep benthics, *Diplotaxodon* have haplotypes that are clearly distinct from those in the rest of the radiation, while the majority of deep benthic haplotypes are their nearest neighbours (Supplementary Fig. 22). The short-wavelength opsin SWS1 is among the genes with high ∆_*N-S*_ scores but it does not exhibit shared selection between *Diplotaxodon* and deep benthics - it is most variable within the shallow benthic group. Finally, the short-wavelength opsins SWS2A and SWS2B have negative ∆_*N-S*_ scores in our Lake Malawi dataset and thus are not among the candidate genes.

There have been many previous studies of selection on opsin genes in fish (e.g. reviewed in ^69–71^), including selection associated with depth preference, but having whole genome coverage allows us to investigate other components of primary visual perception in an unbiased fashion. We found shared patterns of selection between deep benthics and *Diplotaxodon* in the genealogies of six other vision associated candidate genes: a homolog of the homeobox protein *six7*, the G-protein-coupled receptor kinase GRK7, two copies of the retinal cone arrestin-C, the α-subunit of cone transducin GNAT2, and peripherin-2 (Fig. 6a). The functions of these genes suggest a prominent role of cone cell vision in depth adaptation. The homeobox protein *six7* governs the expression of RH2 opsins and is essential for the development of green cones in zebrafish^72^. One of the variants in this gene that distinguishes deep benthic and *Diplotaxodon* is just a residue away from the DNA binding site of the HOX domain, while another is located in the SIX1_SD domain responsible for binding with the transcriptional activation co-factor of *six7*^73^ (Supplementary Fig. 21c). The kinase GRK7 and the retinal cone arrestin-C genes have complementary roles in photoresponse recovery, where arrestin produces the final shutoff of the cone pigment following phosphorylation by GRK7, thus determining the temporal resolution of motion vision^74^. Note that bases near to the C-terminus in RH2Aβ mutated away from serine (S290Y and S292G), thus reducing the number of residues that can be modified by GRK7 (Supplementary Fig. 21b). The transducin subunit GNAT2 is located exclusively in the cone receptors and is a key component of the pathway which converts light stimulus into electrical response in these cells^75^. The final gene, peripherin-2, is essential to the development and renewal of the membrane system in the outer cell segments that hold the opsin pigments in both rod and cone cells^76^. Cichlid green-sensitive opsins are expressed exclusively in double-cone photoreceptors and the wavelength of maximum absorbance in cells expressing a mixture of RH2Aβ with RH2B (*λ*_max_ = 498nm) corresponds to the part of light spectrum that transmits the best into deep water in Lake Malawi^71^. Figure 6c illustrates the possible interactions of all the above genes in a double-cone photoreceptor of the cichlid retina.

Haemoglobin genes in teleost fish are located in two separate chromosomal locations: the minor ‘LA’ cluster and the major ‘MN’ cluster^77^. The region around the LA cluster has been highlighted by selection scans among four *Diplotaxodon* species by Hahn et al.^78^, who also noted the similarity of the haemoglobin subunit beta (HBβ) haplotypes between *Diplotaxodon* and deep benthic species. We confirmed signatures of selection in the two annotated LA cluster haemoglobins. In addition, we found that four haemoglobin subunits (HBβ1, HBβ2, HBα2, HBα3) from the MN cluster are also among the genes with high selection scores (Supplementary Fig. 23). It appears that shared patterns of depth selection may be particular to the β-globin genes (Supplementary Fig. 23b), although this hypothesis must remain tentative due to the highly repetitive nature of the MN cluster limiting our ability to confidently examine variation in all the haemoglobin genes in the region.

A key question concerns the mechanism leading to the similarity of haplotypes in *Diplotaxodon* and deep benthics. Possibilities include parallel selection on variation segregating in both groups due to common ancestry, selection on the gene flow that we described in a previous section, or independent selection on new mutations. From considering the haplotype trees and *f*_*dM*_ statistics summarizing local patterns of excess allele sharing, there is evidence for each of these processes acting on different genes. The haplotype trees for rhodopsin and HBβ have outgroup taxa appearing at multiple locations on their haplotype networks, while *A. calliptera* specimens also appear at divergent positions (Fig. 6a). This suggests that the haplotype diversity of these genes may reflect ancient differences in the founders. In contrast, networks for the green cone genes show patterns more consistent with the Malawi radiation all being derived with respect to outgroups (or with us not having sampled a source of ancestral variation) and we found substantially elevated *f*_*dM*_ scores extending for around 40kb around the RH2 cluster (Fig. 6d), consistent with adaptive introgression in a pattern reminiscent of mimicry loci in *Heliconius* butterflies^79^. In contrast, the peaks in *f*_*dM*_ scores around peripherin-2 and one of the arrestin-C genes are relatively narrow, with boundaries that correspond almost exactly to the gene boundaries. Furthermore, these two genes have elevated *f*_*dM*_ scores only for non-synonymous variants Supplementary Fig. 24, while synonymous variants do not show any excess allele sharing between *Diplotaxodon* and deep benthics. Due to the close proximity of non-synonymous and synonymous sites within the same gene, this suggests that for these two genes there may have been independent selection on the same *de novo* mutations.

## Discussion

Variation in genome sequences forms the substrate for evolution. Here we have described genome variation at the full sequence level across the Lake Malawi haplochromine cichlid radiation. We focused on ecomorphological diversity, representing more than half the genera from each major group, rather than obtaining deep coverage of species within any particular group. Therefore, we have more samples from the morphologically highly diverse benthic lineages than, for example, the mbuna where many species are largely recognised by colour differences.

The observation that cichlids within an African Great Lake radiation are genetically very similar is not new^80^, but we now quantify the relationship of this to within-species variation, and the consequences for variation in local phylogeny across the genome. The observation that between-species divergence is only slightly higher than within species diversity, with cases where sequence divergence within single individual is higher than between species, is likely the result of the young age of the radiation, the relatively low mutation rate, and of the gene flow between taxa. Within-species diversity itself is relatively low for vertebrates, at around 0.1%, suggesting that low genome-wide nucleotide diversity levels do not necessarily limit rapid adaptation and speciation. This conclusion appears in contrast for example with a recent report that high diversity levels may have been important for rapid adaptation in Atlantic killifish^81^. One possibility is that in cichlids repeated selection has maintained diversity in adaptive alleles for a range of traits that support ecological diversification, as we have concluded for rhodopsin and HBβ and appears to be the case for some adaptive variants in sticklebacks^82^.

We provide evidence that gene flow during the radiation, although not ubiquitous, has certainly been extensive. Overall, the numerous violations of the bifurcating species tree model suggest that full resolution of interspecies relationships in this system will require network approaches (see e.g. ref. ^37^; section 6.2) and population genomic analyses within the framework of structured coalescent with gene flow. However, we note that the majority of the signals affect groups of species, suggesting events involving their common ancestors, or are between closely-related species within the major ecological groups. We see only one strong and clear example of recent gene flow between more distantly-related species, not within Lake Malawi itself but between *Otopharynx tetrastigma* from crater Lake Ilamba and local *A. calliptera.* Lake Ilamba is very turbid and the scenario is reminiscent of cichlid admixture in low visibility conditions in Lake Victoria^83^. It is possible that some of the earlier signals of gene flow between lineages we observed in Lake Malawi may have happened during low lake level periods when the water is known to have been more turbid^30^.

Our suggested model of the early stages of radiation in Lake Malawi (Fig. 4f) is broadly consistent with the model of initial separation by major habitat divergence^24^, although we propose a refinement in which there were three relatively closely-spaced separations from a generalist *Astatotilapia* type lineage, initially of pelagic genera *Rhamphochromis* and *Diplotaxodon*, then of shallow- and deep-water benthics and utaka (a clade which includes Kocher’s sand dwellers^22,24^), and finally of mbuna (Fig. 4f). Thus, we suggest that Lake Malawi contains three separate haplochromine cichlid radiations, stemming from the *Astatotilapia calliptera* riverine generalist lineage, and interconnected by subsequent gene flow.

The finding that cichlid-specific gene duplicates do not tend to diverge particularly strongly in coding sequences (Fig. 5b) suggests that other mechanisms of diversification following gene duplications may be more important in cichlid radiations. Divergence via changes in expression patterns has been illustrated and discussed in ref. ^11^. Future studies that address larger scale structural variation between cichlid genomes will be able to assess the contribution of differential retention of duplicated genes.

The evidence concerning shared adaptation of the visual and oxygen-transport systems to deep-water environments between deep benthics and *Diplotaxodon* suggests different evolutionary mechanisms acting on different genes, even within the same cellular system. It will be interesting to see whether the same genes or even specific mutations underlie depth adaptation in Lake Tanganyika, which harbours specialist deep water species in least two different tribes^84^ and has a similar light attenuation profile but a steeper oxygen gradient than Lake Malawi^64^.

Over the last few decades, East African cichlids have emerged as a model for studying rapid vertebrate evolution^11,24^. Taking advantage of recently assembled reference genomes^11^, our data and results provide unprecedented information about patterns of sequence sharing and adaptation across the Lake Malawi radiation, with insights into mechanisms of rapid phenotypic diversification. The data sets we have generated are openly available (see Acknowledgements) and will underpin further studies on specific taxa and molecular systems. For example, we envisage that our results, clarifying the relationships between all the main lineages and many individual species, will facilitate greater use of the Lake Malawi cichlids in speciation studies which require investigation of taxon pairs at varying stages on the speciation continuum^85,86^, and on the role of adaptive gene flow in speciation.

## Methods

The analyses presented above are based on SNPs obtained from Illumina short (100bp-125bp) reads, aligned to the *Metriaclima zebra* reference assembly version 1.1^11^ with `bwa-mem`^87^, followed by GATK haplotype caller^88^ and samtools/bcftools^89^ variant calling, filtering, genotype refinement, imputation, and phasing in BEAGLE^90^ and further haplotype phasing with `shapeit v2`^91^, including the use of phase-informative reads^92^. For details and for descriptions of subsequent analysis methods please see the Supplementary Methods section.

## Acknowledgments

This work was supported by the Wellcome Trust (097677/Z/11/Z to MM, WT098051 to RD and HS), the Royal Society-Leverhulme Trust Africa Awards (AA100023 and AA130107 to MG and GFT) and the European Molecular Biology Organization (ALTF 456-2016 to MM). Raw sequencing reads are in the Sequence Read Archive: (BioProjects PRJEB1254 and PRJEB15289: sample accessions listed in Supplementary Table 4). Whole-genome variant calls in the Variant Call Format (VCF), phylogenetic trees and protein coding sequence alignments are available from the Dryad Digital Repository (http://dx.doi.org/accession). RD declares that he owns stock in Illumina from previous consulting. We want to thank the Sanger Institute sequencing core for DNA sequencing, Mingliu Du for DNA extractions, David Swofford and Michael Matschiner for advice on phylogenomic analyses, Walter Salzburger and Ian Wilson for comments on the manuscript, and the Tanzania Fisheries Research Institute and the Malawi Government Fisheries Research Unit for support.

## Author contributions

EM, GFT, MJG, MM and RD devised the study. GFT and MJG collected the samples. AMT bred parent-offspring trios and performed geometric morphometric analyses. MM performed the DNA extractions. HS and MM analysed the genomic data. All authors participated in interpretation of the results. MM, HS and RD drafted the manuscript, and all others commented.

